# Different artificial lighting spectra change the mating behavior of generalist predator *Orius insidiosus* (Say), and photoperiod extension promotes its development

**DOI:** 10.1101/2025.01.24.634755

**Authors:** Morgane L. Canovas, Paul K. Abram, Roselyne Labbé, Jean-François Cormier, Tigran Galstian, Martine Dorais

## Abstract

In protected cropping systems such as greenhouses and indoor farming, augmentative biological control depends on release rates, establishment, and reproduction of natural enemies. Light-emitting diodes (LEDs) are widely used to enhance plant growth in these systems and are increasingly implemented in mass-rearing facilities for natural enemies. However, the impacts of LEDs on the life cycle of beneficial predators remain insufficiently explored. This study examined the mating behaviors and developmental performance of generalist predator *Orius insidiosus* under light spectra previously shown to support its predation of the pest thrips *Frankliniella occidentalis*. In laboratory experiments, predator pairs were exposed to artificial light sequences starting with a 12h base light condition simulating a cloudy winter day, supplemented by 8h photoperiod extensions (blue, blue-red, or blue-green-red spectra), and a control without extension. Mating occurred under all tested conditions, but blue light reduced mating probability, frequency, and duration. Photoperiod extension improved fecundity, fertility, and second-generation numbers of *O. insidiosus* adults, with blue light favoring egg laying and hatching but not metamorphosis into adults. The second-generation sex ratio was unaffected by light sequence, maintaining population viability with a balanced proportion of females. Our findings demonstrate that *O. insidiosus* can successfully mate, reproduce, and develop under artificial lighting and highlight the potential of modulating light spectrum to optimize both mass-rearing and establishment in protected crops.

**Highlights:**

- Photoperiod extension enhances development of the predatory bug *O. insidiosus*.
- Blue light reduces *O. insidiosus* mating probability, frequency, and duration.
- Photoperiod extension with blue light favors *O. insidiosus* fertility and fecundity.
- *Orius insidiosus* sex ratio is unaffected by photoperiod.
- LEDs could enhance natural enemy establishment in protected crops.

## Introduction

The lighting environment is a crucial driver of insect population dynamics, regulating activity at both behavioral and physiological levels (Johansen et al., 2011; Khyati et al., 2017). Augmentative biological control in protected cropping systems that have a variety of natural and artificial light sources, such as greenhouses and indoor farming, depends on the establishment and reproduction of beneficial insects to ensure continuous pest suppression across multiple generations (Hajek and Eilenberg, 2018). Both inundative and inoculative strategies further require a consistent supply of beneficial insects for repeated introductions throughout the cropping season. In these systems, biological control is increasingly implemented under Light Emitting Diodes (LEDs), which are widely used to enhance plant growth by supplementing natural solar radiation in northern-latitude greenhouses (Dorais et al., 2017) or replacing it entirely in indoor farm settings (Cary and Stutte, 2017). Given their energy efficiency and compact size, LEDs are also increasingly adopted in insect mass-rearing facilities (Ortiz et al., 2016). They can also be programmed to meet the unique lighting needs of different insect species. Arthropods rely on daily changes in natural light wavelengths and photoperiod duration to adjust their biological rhythms (Abrieux et al. 2020) a process mediated by a complex network of photoreceptors at both the organ and molecular levels (Saunders, 2012). In particular, variations in photoperiod and light spectral composition have been shown to regulate the reproductive behaviors and developmental performance of both pest and beneficial insect species (Philogene and McNeil, 1984; Johansen et al., 2011). Thus, manipulating daily light cues through dynamic LED horticultural lighting is likely to influence biological control outcomes and potentially improve mass-rearing efficiency. Although the biological clock mechanism is relatively conserved among arthropods, species-specific differences remain (Abrieux et al. 2020), underscoring the need for further research on key horticultural species.

While greenhouses support year-round plant production, the efficacy of beneficial insects introduced into these environments for biological control often decreases in winter due to the short natural photoperiod. Under light-limiting conditions, many insects enter diapause, an adaptive strategy of physiological dormancy, characterized by arrested feeding and reproductive activities, enabling survival in unfavorable winter environments (Gullan and Cranston, 2014). However, diapause in predatory insects can be prevented by artificially extending the photoperiod (e.g., Berkvens et al., 2008). Exposure to long photoperiods can also improve egg-laying and hatchability (Cochard et al., 2019; Gonzalez et al., 2023) and lead to faster development times (Berkvens et al., 2008; Joschinski et al., 2019; Nissinen et al., 2017), increased prey consumption (Joschinski et al., 2019) or host parasitism rates (Cochard et al., 2019). Accordingly, extended photoperiods are often employed in mass rearing of insects to maximize reproduction and avoid diapause (Abrieux et al. 2020). Compared to high-pressure sodium lamps (HPS), LEDs offer the advantage of light spectrum modulation, significantly influencing the fecundity and sex ratios of biological control agents depending on the predator or parasitoid species (Nissinen et al. 2017; Cochard et al. 2019; Gonzales et al. 2023).

Predators of the *Orius* genus (Hemiptera: Anthocoridae) are key beneficial insects widely used in northern latitudes for augmentative biological control (Van Lenteren, 2012; Van Lenteren et al., 2018). *Orius insidiosus* (Say) is frequently introduced to control various pests commonly found year-round in vegetable and ornamental crops, including the cosmopolitan thrips *Frankliniella occidentalis* (Pergande) (Reitz et al., 2020). The control efficiency of *O. insidiosus* can be reduced from October to March due to the onset of facultative winter reproductive diapause (Van Den Meiracker, 1994). Thus, responses of *Orius* spp. to variable photoperiods have been extensively studied, with findings indicating that at least 10 to 12 hours of daily light exposure effectively alleviates diapause (Bahşi and Tunç, 2012; Herrick et al., 2021; Kingsley and Harrington, 1982; Ruberson et al., 2000; Stack and Drummond, 1997, 1995; Taniai et al., 2021; Van Den Meiracker, 1994). However, the effect of light spectra on other aspects of *Orius* spp. reproductive biology remains underexplored, despite indications that it can influence their reproductive behaviors, fecundity, fertility, and development time (Labbé and McCreary, 2020; Wang et al., 2013). This knowledge gap is particularly relevant for *O. insidiosus*, as its winter establishment remains a recurring challenge for northern greenhouse growers, especially at key periods such as during tomato transplantation in January under artificial lighting, when early preventive releases are critical for effective biological control programs. While *O. insidiosus* maintains predation performance under artificial lighting, its behavior is strongly influenced by light spectra (Canovas et al., 2024). Still, the effects of photoperiod extension with different spectra on its full life cycle, including mating behaviors, remain unknown (Labbé and McCreary, 2020).

In this study, we investigated the effects of artificial light spectra, previously identified as favorable for *O. insidiosus* predation against thrips (Canovas et al., 2024), on the predator’s mating behavior and development. Using a flexible optical device, we examined fine-scale mating behaviors across three spectra. These same spectra were then used to extend a 12-hour base light condition, simulating a cloudy winter day, with 8-hour photoperiod extensions, while monitoring developmental performance. We predicted that extending daylight would enhance reproduction, particularly by promoting egg-laying and hatching under short-wavelength spectra (blue and green) (Cochard et al., 2019b; Labbé and McCreary, 2020; Gonzalez et al., 2023). Conversely, based on findings from another *Orius* species, we anticipated that blue light would suppress mating behaviors, resulting in fewer encounters and shorter mating durations (Wang et al., 2013). This study aimed to evaluate whether photoperiod extension with specific artificial spectra could support *O. insidiosus* reproduction and development, facilitating its establishment for augmentative biological control or mass-rearing applications. Furthermore, we explored whether spectra that enhance predation could also affect population growth (*via* effects on reproduction and development), contributing to the development of strategies to optimize biological control in modern greenhouse systems.

## Material and methods

### Lighting device and treatments

To investigate the effects of light spectrum and photoperiod extension with selected spectra on mating and development of *O. insidiosus*, we used the same laboratory setup described by Canovas et al. (2024). The experiment was performed in the laboratory, in four opaque growth tents (©2022 VIVOSUN, Philadelphia, Ontario; dimensions: L=121.9 cm, W=121.9 cm, H=182.9 cm) outfitted with horticultural dynamic spectrum lighting fixtures (©2022 SOLLUM Technologies, Montreal, Quebec; model SF05-A). These LED fixtures provided real-time control over spectral composition and light intensity via the SUN as a Service® platform (“SUNaaS”). Environmental conditions were maintained at 21 ± 2°C and 60 ± 2% relative humidity using a combination of climate control devices (©2022 Inkbird, Shenzhen, China; models ITC-308 and IHC-200; ©2022 VIVOSUN, Philadelphia, Ontario, CA; model 43237-2; ©2022 ALACRIS, Ottawa, Ontario, CA; 4L humidifier). Each growth tent corresponded to one of the tested light sequences. Based on a previous series of experiments (Canovas et al., 2024), we selected three lighting spectra already characterized as conducive for *O. insidiosus* predation against thrips: 100% Blue (B), 25% Blue – 25% Green – 50% Red (BR), and 10% Blue – 90% Red (BGR). These spectra were used to supplement an artificially recreated natural 12-hour photoperiod of low-light conditions, followed by an 8-hour extension, simulating winter greenhouse lighting in northern latitudes (Fig. 1). Variations in light intensity and spectral composition over a 12-hour cycle of “natural” light were based on average spectrometric data provided by the World Meteorological Organization (WMO) for the Quebec City region and implemented in the SUNaaS platform (Honnorine Lefevre, agr., personal communication; SOLLUM Technologies). A control treatment (CTRL), without any photoperiod extension, was also included in the experimental design. To facilitate manipulations, the “natural” 12-hour light phase occurred from 7 PM to 7 AM, followed by the extension phase from 7 AM to 3 PM and a 4-hour period of darkness, except for the control treatment (12 hours of darkness from 7 AM to 7 PM) (Fig. 1). Light intensities in experiments were standardized at 240 µmol/m²/s during the primary photoperiod and approximately 30 µmol/m²/s for the photoperiod extension, a range of intensities over which *O. insidiosus* predation behavior did not change in our prior study (Canovas et al. 2024). Light sequences were randomly reassigned to a new opaque tent at the start of each temporal repetition. Since a photoperiod of at least 12 hours was provided, and *O. insidiosus* neither enters diapause nor reduces its predation activity under such conditions in laboratory or greenhouse settings (Van Den Meiracker, 1994; Herrick et al., 2021), this setup enabled the comparison of the effects of photoperiod extension and the spectrum used for this extension on the reproductive and developmental performance of *O. insidiosus* without triggering diapause.

**Fig. 1.**
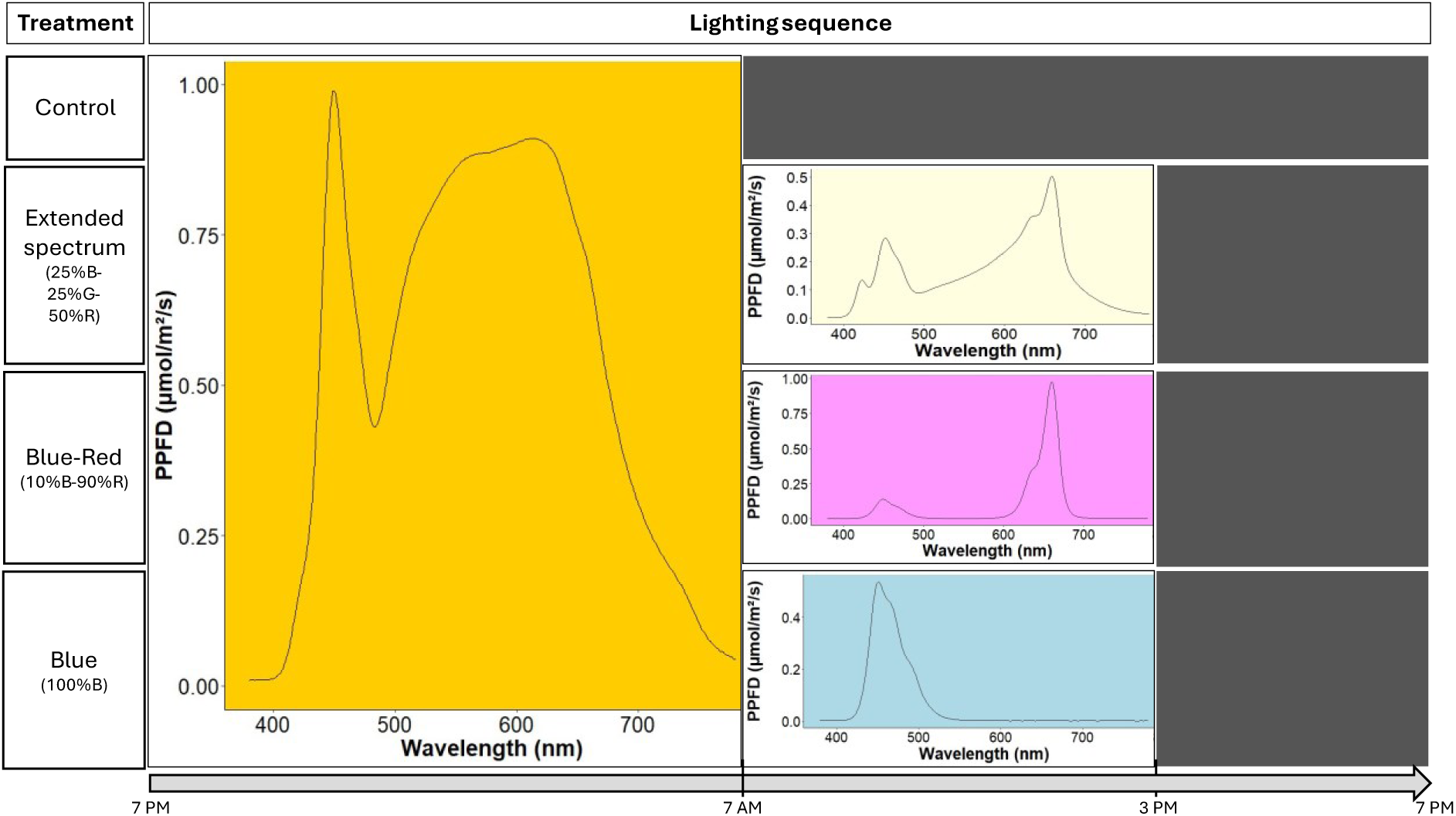
Lighting sequences tested in growth tents during both the mating behavior and the daily light supplementation experiments. The 24-hour cycle was divided into three parts: an initial phase of exposure to an artificially recreated ‘solar’ spectrum (indicated in orange), followed by a photoperiod extension phase using one of the three tested light spectra (indicated in either beige (BGR), pink (BR) or blue (B); except for the control treatment (CTRL)), and finally, a dark phase (indicated by black rectangles). In each treatment, the solar spectrum was artificially recreated, including a sunrise and sunset phase (ʎ_max_= 560 nm). Observations for the mating behavior experiment were realized during the extension phase. The light environment was characterized using a LI-180 Spectrometer (LI-COR, Tucson, Arizona, USA), measured at Petri dish height

### Mating behavior

To compare the impact of lighting spectra treatments (B, BR, BGR, CTRL) on small scale mating behaviors, we purchased age-synchronized *O. insidiosus* from a biological control supplier (©ANATIS Bioprotection, Saint-Jacques-le-Mineur, Quebec, CA) for periodic rearing during the trials. Upon receipt, adults *O. insidiosus* were placed in transparent 1 L Mason jars with the lid replaced by Nitex (160 µm nylon mesh screens). Each jar contained approximately 3-4 cm of a mixture of buckwheat husks and vermiculite, as well as a piece of absorbent paper. Six commercial green bean pods, disinfected for 10 minutes in a 5% dilution of bleach solution (LAVO PRO 6), provided a water source and oviposition substrate. Six 2 cm-diameter stickers coated with frozen *Ephestia kuehniella* eggs (©ANATIS Bioprotection) served as the primary food source (Labbé et al., 2018). The green beans and *E. kuehniella* eggs were replaced every 3 days. A Mason jar, serving as a rearing unit, was placed in each of the four growth tents described in the “Lighting Device and Treatments” section, with each tent corresponding to one of the tested light sequences. Unlike previous studies on *Orius* spp. reproduction (Leon-Beck and Coll, 2009; Arakawa et al., 2019; Wang et al., 2013; Vacacela Ajila et al., 2019; Taniai et al., 2021; Rodríguez-Gómez et al., 2022), we chose to use male and female predators sourced from a commercial shipment with a typical 45% female:55% male ratio, rather than laboratory-reared naïve individuals. This approach was designed for realism, using predators sourced from a supplier common to commercial greenhouse and indoor farming. Males and females were in contact during commercial rearing, transport, and temporary laboratory rearing, allowing mating to occur before experiments. While a single mating is sometimes sufficient for Heteroptera, additional matings can occur (Lundgren, 2011), including in Anthocoridae (Saulich and Musolin, 2009). Previously mated *Orius insidiosus* females subjected to storage, handling, and shipment exhibit higher fecundity than virgins (Bueno et al., 2014). While monoandry has been suggested for *O. insidiosus*, consecutive matings are possible (Vacacela Ajila et al., 2019), and our results show high female receptivity, with an average mating probability of 58% ± 15%.

On the day of the experiments, predators from each lighting sequence treatment were individually placed in transparent 1.5 ml microtubes (©Sarstedt) for sex determination under a stereomicroscope using morphological criteria (Isenhour and Yeargan, 1981). Each *O. insidiosus* pair was then exposed to 30 minutes of darkness, after being placed in their original opaque tent for a 1-minute acclimation to the tested light treatment, ensuring artificial lighting effects were isolated from laboratory ambient light conditions (Cochard et al., 2017). The male and female were subsequently introduced into a sterile 3.5 cm Petri dish (©Fisherbrand™), and their behaviors were recorded for 15 minutes (1080p, 30 fps). Each *O. insidiosus* pair was observed only once. In the opaque tents used for the experiment, only 2% of incoming light was reflected by the Petri dish lid. Observations were conducted between 8:00 AM and 2:00 PM, during the photoperiod extension phase for all treatments except the control (Fig. 1). Six predator pairs were observed simultaneously each day for each of the four tested spectra, over two consecutive days. The experiment was repeated twice in time with separate batches of predators to account for variation in vigor between supplier shipments, resulting in 24 replicates per spectrum treatment.

The video recordings of these trials were analyzed using the open-source Behavioral Observation Research Interactive Software (BORIS) (Friard and Gamba, 2016). Mating frequency and duration are critical factors influencing the fecundity of female predatory Heteroptera, with multiple matings often resulting in increased fecundity (Lundgren, 2011). However, the only previous study examining mating behavior in an *Orius* species under artificial lighting relied solely on direct observations, offering limited insights beyond mating duration (Wang et al., 2013). To address this gap, we estimated the following parameters: (i) mating probability (the likelihood of at least one mating event occurring); (ii) total number of mating events; and (iii) mean duration of mating events within the 15-minute observation window. A mating event was defined as the male’s abdomen contacting the female’s body for more than 3 seconds, based on the minimum duration reported for an *Orius* sp. (Askari and Stern, 1972). This threshold was chosen due to the variability in mean mating durations observed across *Orius* species and experimental conditions (Askari and Stern, 1972; Wang et al., 2013; Arakawa et al., 2019; Vacacela Ajila et al., 2019; Rodríguez-Gómez et al., 2022).

### Development under daily light supplementation

To compare the impact of photoperiod extension, as well as extension spectrum, on *O. insidosus* development, new predator couples originating from the same rearing units as for the mating experiment were placed in experimental units. Each unit consisted of a sterile 9 cm Petri dish (©Fisherbrand™), with a modified lid including a 4 cm circle covered with Nitex for ventilation. A Whatman paper disk (©Fisher Scientific) was placed at the bottom of each Petri dish, along with half a dental cotton roll saturated with water to maintain humidity. A green bean segment approximately 3 cm in length, previously disinfected with bleach and sealed with food-grade wax, served as the oviposition substrate. Additionally, half of a 2 cm diameter sticker coated with frozen *E. kuehniella* eggs was included to standardize resource availability. All units were sealed with medical tape, with 10 units per light sequence tested for each temporal repetition. The male and female predators were kept together for 48 hours, after which the male was removed. Over the following six days, the number of eggs laid by the female during each 48-hour period was recorded, resulting in 60 observations per light sequence tested. Green bean segments and *E. kuehniella* egg sources were systematically replaced every two days. After egg counts were completed, each green bean segment was transferred to a 500 ml Mason jar with lid replaced by Nitex, containing a 5 cm Whatman paper disk and a dental cotton roll saturated with water. These jars were then returned to the corresponding parental light sequence. Over time, green bean segments from the same experimental unit were combined within the same jar, resulting in 10 jars per tested light sequence. Two bleach-disinfected green bean pods and two 2 cm diameter stickers coated with frozen *E. kuehniella* eggs were added to each jar two days after the first segment was incubated and were subsequently replaced every two days.

Ten days after the start of the experiment, the total number of juveniles, across all developmental stages, was recorded. Juveniles were then transferred to a 500 ml Mason jar containing Whatman filter paper, a damp dental cotton roll, two pods and two stickers coated with frozen eggs being replaced every two days as previously described. The 10-day period for juvenile counts was chosen based on Isenhour and Yeargan (1981), who reported an average hatching time of 6.5 days for *O. insidiosus* eggs at 20–24°C, ensuring most eggs had hatched. This timing also minimizes overlap with later developmental stages while optimizing data collection efficiency. The jars were inspected daily for metamorphosis into adults (i.e. emergence of adult individuals). The number of adults was recorded daily, and adults were immediately transferred to 70% alcohol for sex determination under a stereomicroscope (Isenhour and Yeargan, 1981). This process was repeated until no further adult emergence was observed. The mating experiment was repeated twice with separate batches of predators to account for variation in vigor between supplier shipments

### Statistical analysis

The data were analyzed using R software (R Core Team, 2024; version 4.4.1), with a statistical significance threshold set at p = 0.05. All response variables were analyzed using generalized linear mixed models (GLMMs) fitted with the `glmmTMB` function from the `glmmTMB` package (Brooks et al., 2017). Temporal repetitions with distinct insect batches were specified as a random effect to account for variability across experimental runs. For the mating behavior experiment, the following distributions and link functions were specified: a Binomial error distribution (logit link function) for mating probability, a Tweedie error distribution (log link function) for mean mating event duration, and a Poisson error distribution (log link function) for the number of mating events. In the daily light supplementation experiment, egg laying, the number of juveniles, and the number of adults were analyzed using a Tweedie error distribution (log link function), while Beta error distributions (logit link function) were used for egg hatching and metamorphosis into adults’ probabilities. The proportion of female insects within the second generation was analyzed with Beta binomial error distribution (logit link function). Pairwise comparisons to distinguish between levels of significant factors were conducted using estimated marginal means (EMMs) with a Tukey adjustment for multiple comparisons (Graves et al., 2024; Lenth, 2024).

In our models, p-values were derived from asymptotic Wald tests, assuming infinite denominator degrees of freedom. Residual diagnostics were performed using the `DHARMa` package (Hartig, 2022), with simulated residuals visually inspected *via* diagnostic plots to assess deviations from model assumptions such as residual uniformity, overdispersion, and zero inflation. Graphical representations illustrate data adjusted from the full model, based on estimated marginal means (Lenth, 2024). All models followed a generalized completely randomized block design, with temporal repetitions treated as blocks (random effects) to account for within-block variability, and each treatment repeated within the same block.

## Results

### Mating behavior

While the mating probability of *O. insidiosus* couples was not affected by light sequence (Table 1), there was a trend towards reduced mating probability (−42%) under blue compared to blue-red light (Fig. 2A). The mean duration of a mating event varied across light sequences (Table 1), with a 71% shorter duration observed under blue light compared to control conditions (Fig. 2B). Similarly, the mean number of mating events was affected by light sequence (Table 1), with 42% fewer events under blue light compared to control (Fig. 2C).

**Fig. 2.**
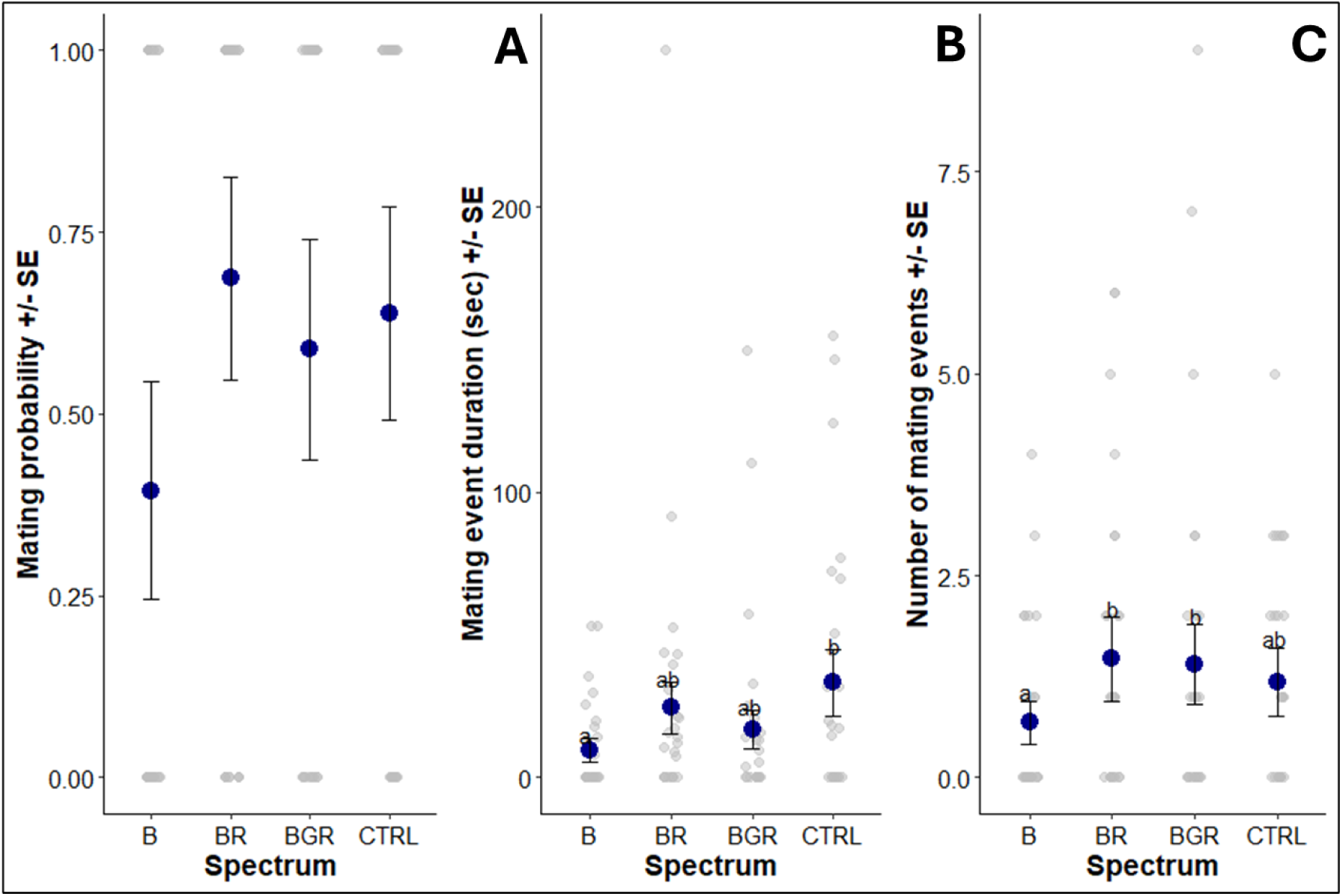
Behavioral responses of *Orius insidiosus* couples under different light spectra: A) Mating probability (± SE); B) Mating event duration (± SE); C) Number of mating events (± SE). The raw data are shown in gray, while the model-predicted means are in blue.

**Table 1.** Statistical comparison of mating behaviors of *Orius insidiosus* couples exposed to different light spectra.

| Measurement | F-value<sub>df</sub> | P-value |
| --- | --- | --- |
| Mating probability<sup>a</sup> | $F_{(3, \infty)} = 1.325$ | 0.264 |
| Mating event duration<sup>b</sup> | $F_{(3, \infty)} = 1.050$ | 0.013 |
| Number of mating events<sup>c</sup> | $F_{(3, \infty)} = 2.876$ | 0.035 |

a: analyzed using generalized linear mixed model with Binomial error distribution
b: analyzed using generalized linear mixed model with Tweedie error distribution
c: analyzed using generalized linear mixed model with Poisson error distribution

### Development under daily light supplementation

The number of eggs laid in 48 hours varied across light sequences (Table 2), with a 66% fewer eggs laid under control (short-day) conditions compared to blue light photoperiod extension (Fig. 3A). Egg hatching probability was also affected by light sequence (Table 2), with a 52% lower probability under control conditions compared to blue and blue-red light treatments (Fig. 3B). The number of juveniles varied according to light sequence (Table 2), with 57% fewer nymphs observed under control conditions compared to blue light (Fig. 4A). Similarly, the metamorphosis into adults’ probability varied across light sequences, showing a 28% reduction under blue light compared to control (Fig. 4B). Although the number of adults was not significantly affected by the light sequence (Table 2), 37% fewer adults were produced under control conditions compared to blue light photoperiod extension (Fig. 5A). Finally, the sex ratio (proportion females) in the second generation was consistent across light sequences (Table 2), with a mean predicted value of 51% ± 2% (Fig. 5B).

**Fig. 3.**
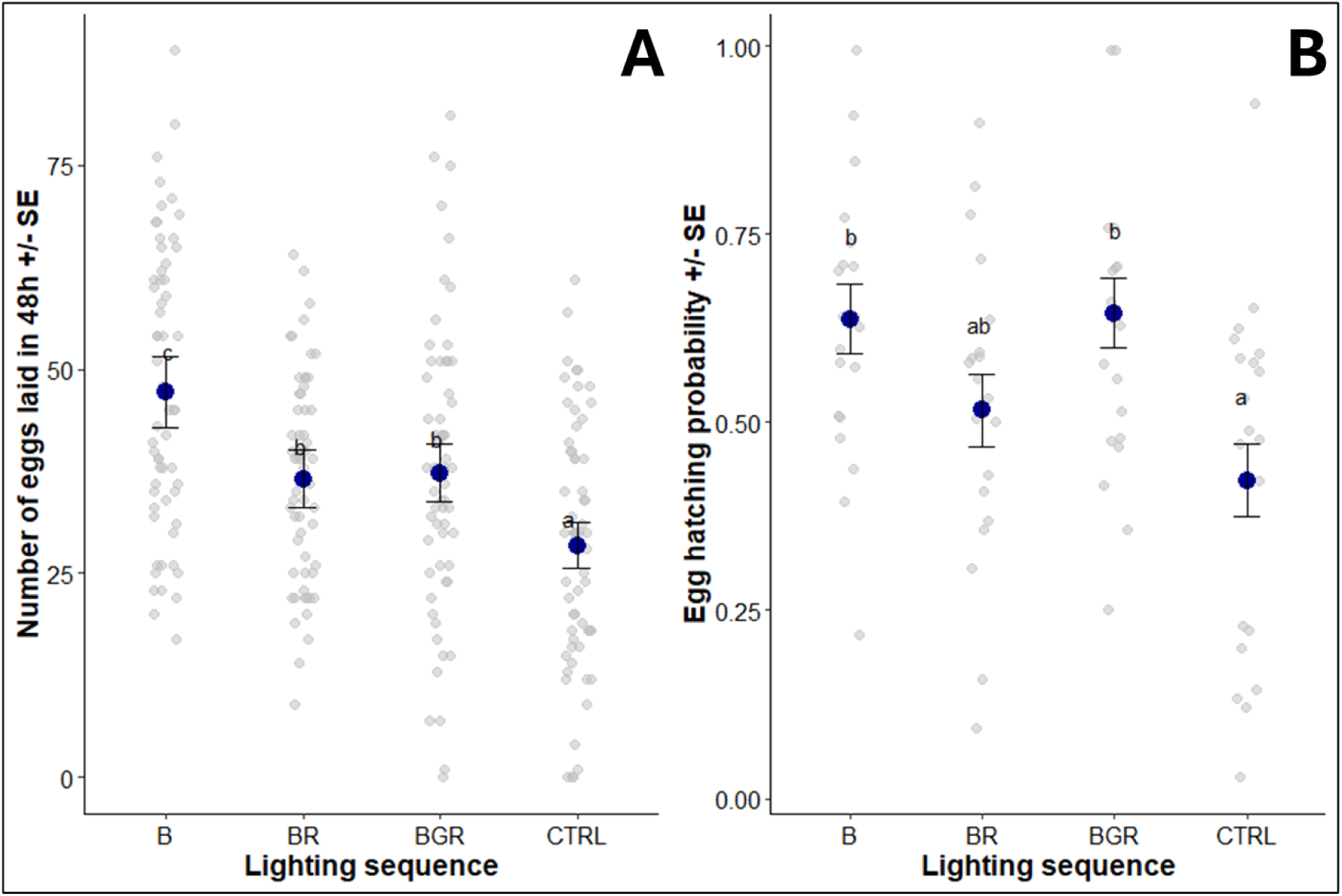
Developmental performance of second-generation of *Orius insidiosus* under different lighting sequences: A) Number of eggs laid in 48h (± SE); B) Egg hatching probability (± SE). The raw data are shown in gray, while the model-predicted means are in blue.

**Fig. 4.**
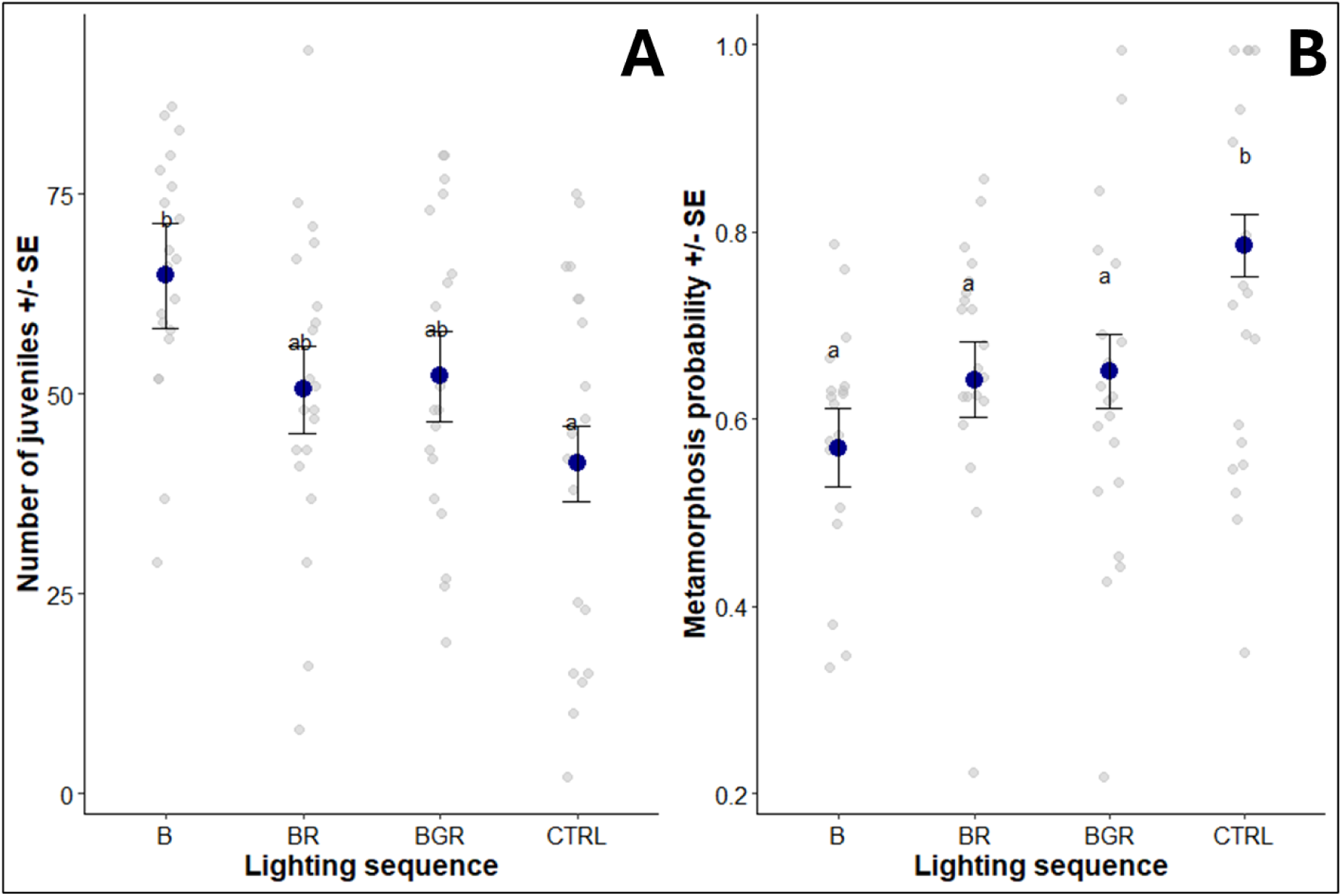
Developmental performance of second-generation of *Orius insidiosus* under different lighting sequences: A) Number of juveniles (± SE); B) Metamorphosis probability (± SE). The raw data are shown in gray, while the model-predicted means are in blue.

**Fig. 5.**
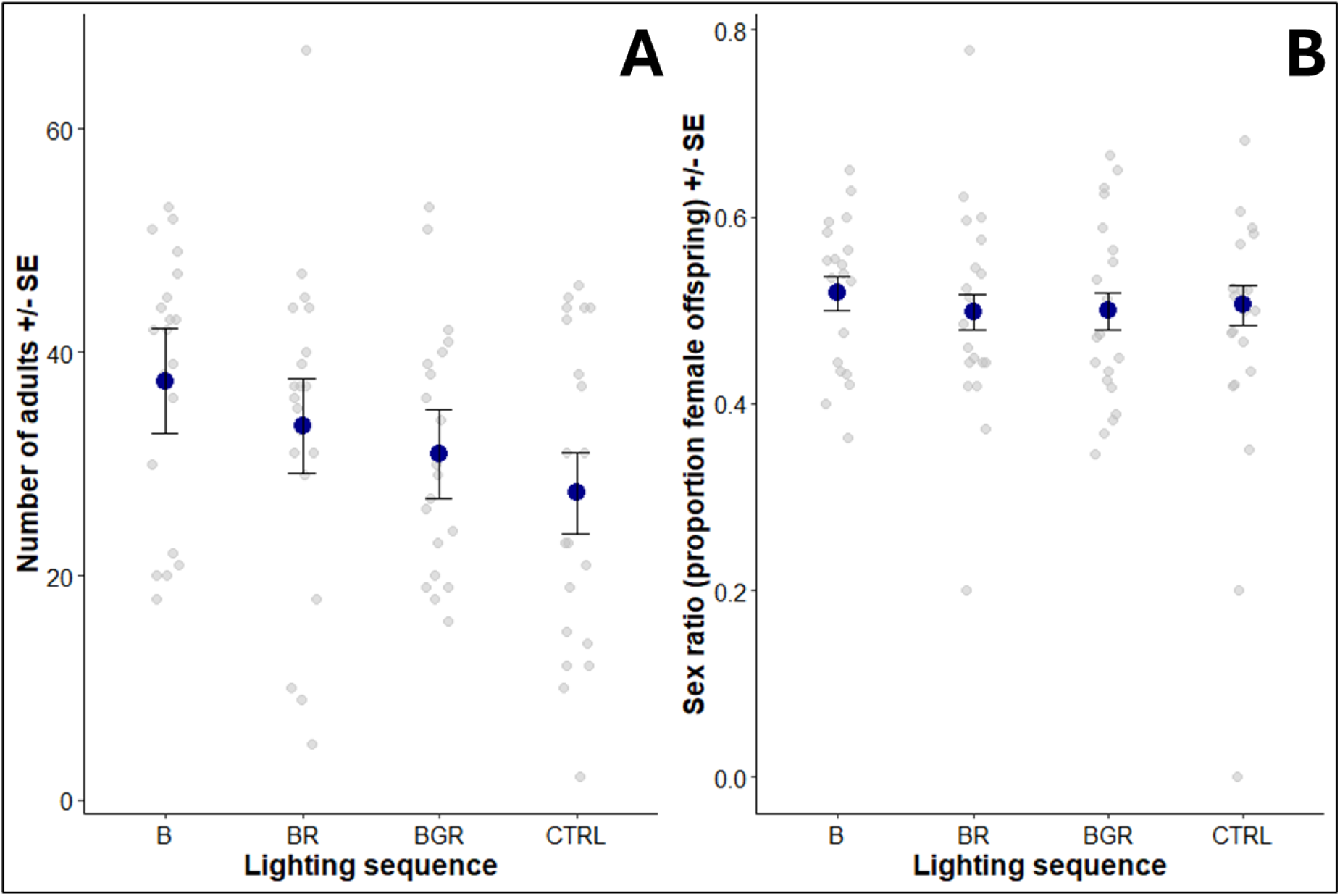
Developmental performance of second-generation of *Orius insidiosus* under different lighting sequences: A) Number of adults (± SE); B) Sex ratio (proportion female offspring) (± SE). The raw data are shown in gray, while the model-predicted means are in blue.

**Table 2.** Statistical comparison of *Orius insidiosus* development parameters while exposed to different photoperiod extension spectra.

| Measurement | F-value<sub>df</sub> | P-value |
| --- | --- | --- |
| Number of eggs laid in 48h<sup>a</sup> | F<sub>(3, ∞)</sub> = 13.569 | < 0.0001 |
| Egg hatching probability<sup>b</sup> | F<sub>(3, ∞)</sub> = 4.722 | 0.003 |
| Number of juveniles<sup>a</sup> | F<sub>(3, ∞)</sub> = 4.615 | 0.003 |
| Metamorphosis probability<sup>b</sup> | F<sub>(3, ∞)</sub> = 5.363 | 0.001 |
| Number of adults<sup>a</sup> | F<sub>(3, ∞)</sub> = 2.289 | 0.076 |
| Proportion of female offspring<sup>c</sup> | F<sub>(3, ∞)</sub> = 0.232 | 0.874 |

a: analyzed using generalized linear mixed model with Tweedie error distribution
b: analyzed using generalized linear mixed model with Beta error distribution
c: analyzed using generalized linear mixed model with Beta binomial error distribution

## Discussion

*Orius insidiosus* successfully mated and developed under all tested light conditions, with the second generation maintaining a balanced sex ratio under sole-source artificial lighting across spectra. Overall, photoperiod extension significantly enhanced the predator developmental performance throughout its life cycle compared to no extension, with blue light having the most beneficial effects. Interestingly, despite reduced mating activity under blue light, this spectrum yielded the highest developmental success, highlighting a complex relationship between reproductive behaviors and life cycle outcomes.

We found that *O. insidiosus* mated under all tested spectra, but blue light significantly reduced mating probability, frequency, and duration, as predicted. This result aligns with a previous study on *Orius sauteri* (Poppius), where both the likelihood and duration of mating were reduced under blue light (Wang et al., 2013). Similarly, blue light was associated with the lowest prey capture performance of *O. insidiosus* (Canovas et al., 2024), suggesting that this wavelength may impair visual cues critical for small-scale interactions with both prey and mating partners. Interestingly, the number of mating events observed in our study was higher than previously reported for *Orius* species under laboratory conditions (Askari and Stern, 1972; Vacacela Ajila et al., 2019). To the best of our knowledge, this is the first study to examine such fine-scale mating behavior under artificial light spectra in an *Orius* species. Understanding the impact of light spectra on reproductive behaviors is crucial for both augmentative biological control and mass-rearing protocols, as suboptimal mating often limits the establishment of beneficial populations in cropping systems (Hopper and Roush, 1993; Queffelec et al., 2021).

Photoperiod extension with artificial lighting enhanced predator developmental performance. As predicted, extended photoperiod under blue light was associated with increased fecundity, fertility, and the highest number of offspring, supporting previous findings on the fecundity of *O. insidiosus* under low-intensity blue light (Labbé and McCreary, 2020). Furthermore, blue light specifically favored egg-laying and hatching but not metamorphosis, confirming that the effects of blue light on insect life cycles are highly species-dependent (Hori et al., 2014) and also vary by life stage (Shibuya et al., 2018). Sex ratios in the second generation were not affected by any of the tested lighting sequences. This result contrasts with previous concerns about skewed sex ratios of some biological control organisms (aphid parasitoids) under red-light-enriched environments (Cochard et al., 2019). However, it concurs with studies on more closely related predatory species; in one such study, photoperiod duration did not affect the sex ratio of *Orius majusculus* (Reuter) (Bahşi and Tunç, 2012). Across all tested treatments, the proportion of female *O. insidiosus* remained above the critical threshold of 45%, as targeted by the biological control supplier (personal communication, Dr. Silvia Todorova, ANATIS Bioprotection). In addition to promoting population growth, maintaining sex ratio stability under artificial lighting is essential for long-term population establishment, reinforcing the utility of optimized lighting in augmentative biological control strategies. *O. insidiosus* appears to be a particularly adaptable beneficial species, maintaining both its predation (Canovas et al., 2024) and developmental capacities (current study) under artificial lighting treatments representative of what is used in horticultural production.

Despite reduced mating of *O. insidiosus* under blue light, we observed higher fecundity, fertility, and number of adults in the second generation exposed to this spectrum. We hypothesize that mating stress underlies this result: spectra that suppress mating activity may promote reproduction. A similar trend was reported for *O. sauteri*, where higher light intensities, associated with shorter mating durations, increased female fecundity (Wang et al., 2013). Traumatic insemination (TI), a reproductive strategy in which the male pierces the female’s body to deposit sperm, has been documented in the Anthocoridae family (Tatarnic et al., 2014; Vacacela Ajila et al., 2019) and likely occurs in *O. insidiosus* (Vacacela Ajila et al., 2019). While sometimes necessary for reproduction, TI can negatively affect female longevity and fertility through energy depletion, physical injury, or infection (Reinhardt et al., 2015), potentially explaining the significantly reduced egg-laying observed under other spectra compared to blue light. Vacacela Ajila et al. (2019) reported that the genital morphology of *O. insidiosus* females provides little protection against TI-related injuries, including occasional fatal damage during mating. Similarly, in *O. sauteri*, multiple matings do not enhance fecundity or hatching success (Arakawa et al., 2019). On the other hand, a male *Orius laevigatus* (Fieber) can effectively fertilize up to six consecutive females (Rodríguez-Gómez et al., 2022), and sexual harassment by males reduces female fecundity (Leon-Beck and Coll, 2009). These findings suggest that light spectrum modulation could indirectly enhance predator population growth by reducing mating frequency and thus mating-related stress.

Our results emphasize the benefits of photoperiod extension with specific spectra in supporting natural enemy establishment through mating and development, as observed in other beneficial insect species. However, the optimal light spectrum for *O. insidiosus* varies depending on the context and life stage. For mass-rearing, a photoperiod extension with blue light is recommended during early development to reduce mating stress and enhance egg-laying and hatching. This should be followed by a transition to broad-spectrum light resembling sunlight to promote adult metamorphosis. In greenhouse settings, a 10% blue-90% red light extension at the end of the day could enhance predation (Canovas et al. 2024), aiding in pest outbreak prevention during at-risk periods. Additionally, photoperiod extension with blue light combined with natural solar radiation is likely to promote *O. insidiosus* growth and establishment after predator introduction in greenhouses. These recommendations require validation under commercial conditions, where other factors like temperature and humidity as well as environmental complexity may also influence biological control outcomes and predator development. While the current study did not investigate the indirect effects of artificial lighting through plant-mediated pathways—which modulate plant-insect interactions (Vänninen et al., 2010; Lazzarin et al., 2021; Meijer et al., 2024, 2023; Zhu et al., 2024)—it is part of a series of studies that will address these aspects. The combined direct and indirect effects of lighting conditions on thrips biological control outcomes with *O. insidiosus* warrant further investigation (Canovas et al., in preparation), aiming to optimize LED-lighting strategies by integrating insect biology and plant-insect interactions within protected cropping systems.

## Declaration of generative AI and AI-assisted technologies in the writing process

During the preparation of this work the authors used ChatGPT-4o for optimizing code implementation and troubleshooting during statistical analyses conducted in RStudio, as well as assisting in the drafting and refinement of the manuscript. After using this tool, the authors reviewed and edited the content as needed and took full responsibility for the content of the published article.

## Acknowledgments

The authors would like to acknowledge the partnership and financial support received from the Ministère de l’Agriculture, des Pêcheries et de l’Alimentation du Québec, MITACS, SOLLUM Technologies inc., ANATIS Bioprotection and the National Optics Institute (project n° IA120649 & IT17200).

## Authors’ contribution

MLC: Conceptualization; Funding acquisition; Project administration; Methodology; Investigation; Formal analysis; Visualization; Writing – original draft; Writing – review & editing. PKA: Supervision; Writing – original draft; Writing – review & editing. RL: Writing – review & editing. JFC: Funding acquisition; Supervision; Writing – review & editing. TG: Funding acquisition; Supervision; Methodology; Writing – review & editing. MD: Funding acquisition; Project administration; Supervision; Resources; Writing – review & editing.

## Ethical approval

The authors approve the ethical guidelines of the journal.

## Consent for publication

All authors gave consent to the publication of the study.

